# Unliganded Progesterone Receptor Governs Estrogen Receptor Gene Expression by Regulating DNA Methylation in Breast Cancer Cells

**DOI:** 10.1101/192567

**Authors:** Gaetano Verde, Lara I. De Llobet, Roni H.G. Wright, Javier Quilez, Sandra Peiró, François Le Dily, Miguel Beato

**Affiliations:** Centre de Regulació Genòmica (CRG), Barcelona Institute for Science and Technology, Barcelona, 08003 Spain, (L.I.D.L.); (R.H.G.W.); (J.O.); (F.L.D.); Vall d’Hebron Institute of Oncology, Barcelona, 08035, Spain; Universitat Pompeu Fabra, Barcelona, 08003 Spain

**Keywords:** progesterone receptor, estrogen receptor, gene expression, DNA methylation, breast cancer, endocrine therapy

## Abstract

Breast cancer prognosis and response to endocrine therapy strongly depends on the expression of the estrogen and progesterone receptors (ER and PR, respectively). Although much is known about ER*α* gene (*ESR1*) regulation after hormonal stimulation, how it is regulated in hormone-free condition is not fully understood. We used ER-/PR-positive breast cancer cells to investigate the role of PR in *ESR1* regulation in the absence of hormones. We show that PR binds to the low-methylated *ESR1* promoter and maintains both gene expression and DNA methylation of the ESR1 locus in hormone-deprived breast cancer cells. Depletion of PR reduces *ESR1* expression, with a concomitant increase in gene promoter methylation. The high amount of methylation in the *ESR1* promoter of PR-depleted cells persists after the stable re-expression of PR and inhibits PR binding to this genomic region. As a consequence, the rescue of PR expression in PR-depleted cells is insufficient to restore *ESR1* expression. Consistently, DNA methylation impedes PR binding to consensus progesterone responsive elements. These findings contribute to understanding the complex crosstalk between PR and ER and suggest that the analysis of *ESR1* promoter methylation in breast cancer cells can help to design more appropriate targeted therapies for breast cancer patients.

## 1. Introduction

Estrogen and progesterone are the main players in the progression and outcome of breast cancers [1]. Both hormones act through their cognate receptors, estrogen receptor (ER) and progesterone receptor (PR) [1]. Upon hormone exposure, PR and ER exhibit enhanced binding to specific DNA sequences known as hormone-responsive elements, which are generally located within target gene enhancers or promoters [1,2]. The DNA-bound receptors orchestrate the assembly of large cofactor-containing protein complexes that can either positively or negatively affect gene transcription [1]. In addition, hormone-activated ER and PR attached to the cell membrane can trigger rapid signaling by interacting with several kinases, which also participate in hormonal gene regulation [1,3].

PR and ER profoundly affect the breast cancer cell biology. Approximately one-third of breast cancers lack both hormone receptors (ER−/PR−) and generally shows poor histological differentiation with higher growth rates [4]. These cancers rarely respond to hormone therapies and exhibit a poor clinical outcome compared to breast cancers that express both hormone receptors (ER+/PR+) [4,5]. In addition, ER-positive tumors lacking PR (ER+/PR−) are less likely to respond to endocrine therapy compared to ER+/PR+ cancers [4,5]. Consistent with this data, unliganded PR enhances breast cancer cell response to estrogen and to the selective ER modulator used for endocrine therapy, including tamoxifen and others antiestrogens [6,7]. However, how the presence of hormone-free PR affects the breast cancer sensitivity to these external stimuli is not completely understood.

About two-thirds of breast cancers are resistant to steroid hormones at the time of diagnosis, but often retain functional hormone receptors and remain highly sensitive to growth factors. Growth factors induce breast cancer cell proliferation by enhancing the transcriptional activity of the hormone-free PR via the protein kinases’ signaling pathways [8]. The phosphorylated and under-SUMOylated unliganded PR positively regulates the expression of growth-promoting genes by recruiting steroid receptor coactivator 1 (SRC1) to gene promoters [9]. In contrast, SUMOylated PR downregulates the expression of the same class of genes by recruiting histone deacetylase 3 (HDAC3) to promoters [9]. Thus, unliganded PR affects breast cancer progression by regulating gene expression at the chromatin level, even if the molecular mechanisms put into play are not completely clear.

The key role of unliganded PR for the responsiveness of breast cancers to endocrine therapy and growth factors [7,8], as well as the high correlation between PR and ER levels in breast cancers [10] led us to explore whether and how PR regulates the ERα gene (*ESR1*) expression in hormone-free breast cancer cells. Here, we report that PR binds to the low-methylated *ESR1* promoter to maintain its basal expression and its low level of DNA methylation in hormone-free breast cancer cells. Consistent with these data, we show that DNA methylation hinders PR binding to hormone-responsive elements

## 2. Results

### 2.1. Unliganded PR is Required to Maintain ESR1 Gene Basal Expression in Breast Cancer Cells

Comparing the *ESR1* gene expression in T47D breast cancer cells and a derived clone selected for its low PR expression (T47D-Y) [11], as previously observed [12], we confirmed that low PR expression is accompanied by a low expression of *ESR1* at both the transcript and protein levels (Figure 1a,b, left panels). Although the PR inhibits *ESR1* gene expression in T47D cells upon progestin exposure [13], the decrease of *ESR1* expression in T47D-Y cells led us to hypothesize that the hormone-free PR could be involved in maintaining basal *ESR1* gene expression. To test this possibility, we knocked down PR in T47D breast cancer cells using the short-hairpin RNA approach (shPR) and analyzed *PR* and *ESR1* gene expression by RT-qPCR and Western blotting assays. Concomitant with the decrease of PR levels, RNA and protein amount of *ESR1* decreased in shPR cells compared to control cells (shC) (Figure 1a,b, right panels; Figure S1a,b). Interestingly, the strongest PR knockdown cells showed the lowest *ESR1* levels (Figure S1a,b), demonstrating a quantitative relationship between the levels of the two hormone receptors.

**Figure 1.**
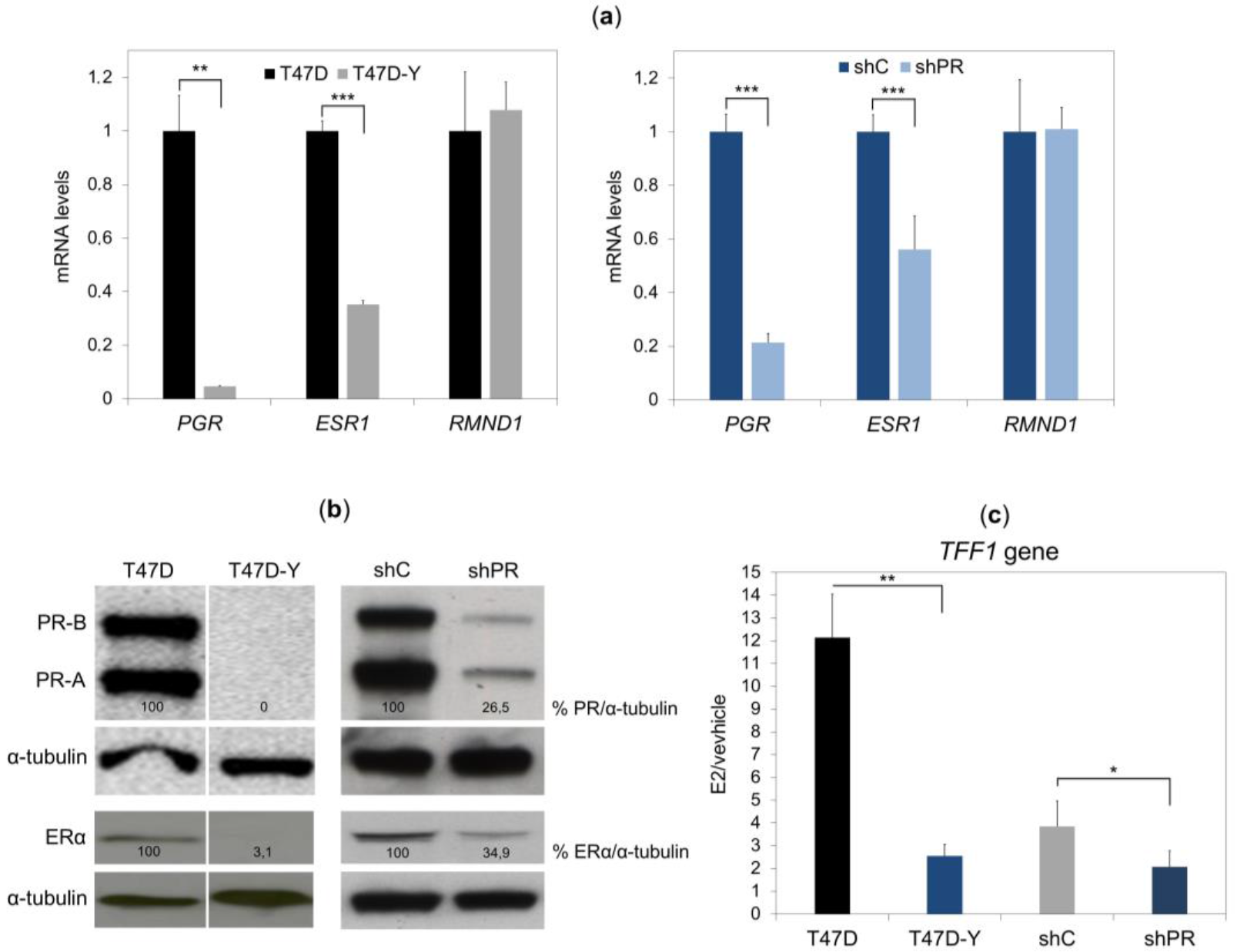
Loss of PR reduces the *ESR1* expression in hormone-deprived T47D breast cancer cells. (**a**) Gene-specific mRNA expression measured by quantitative RT-PCR in T47D or PR-deficient cells (T47D-Y) (left panel) and T47D cells transduced with shRNA against PR (shPR, clone trcn0000010776) or scramble shRNA (shC) (right panel). The gene-specific expression levels were normalized to *GAPDH* expression and are represented as relative values in the T47D cells. *RMND1* was used as a PR-independent control. *PGR*, PR-encoding gene; *ESR1*, ER-encoding gene. Error bars represent the SD of three independent experiments. ** *p* less than or equal to 0.01, *** *p* less than or equal to 0.005, unpaired two-tailed Student’s *t*-test. (**b**) PR and ERα protein levels measured by Western blot in T47D and T47D-Y cells (left panel) and in T47D transduced with shRNA against PR (shPR; clone trcn0000010776) or scramble shRNA (shC) (right panel). α-tubulin protein was used as the loading control. The intensities of the PR and ER bands were normalized to α-tubulin and represented as the relative value in the control cells. The vertical white line depicts a removed lane between the two samples. Blots are representative of three independent experiments. (**c**) PR depletion impairs *TFF1* estrogen-mediated gene transcription. T47D cells, PR-deficient (T47D-Y) cells, short hairpin control (shC) and PR-depleted cells (shPR, clone trcn0000010776) were treated with estradiol (E2, 10 nM) or ethanol (vehicle) for 6 h, at which point *TFF1* mRNA expression was measured by quantitative RT-PCR. The *TFF1* gene expression was normalized to *GAPDH* expression and is represented as fold change relative to the vehicle (E2/vehicle). Error bars represent the standard deviation (SD) of three independent experiments. * *p* less than or equal to 0.05, ** *p* less than or equal to 0.01, unpaired twotailed Student’s *t*-test.

To further confirm the decrease of PR and ER levels in PR-depleted cells (shPR) and PR-deficient cells (T47D-Y), we analyzed the transcription of PR-target or ER-target genes in control cells, T47D-Y and shPR cells treated with progestin (R5020), estradiol (E2) or vehicle (ethanol). The results showed that the progestin-mediated gene expression was strongly affected in T47D-Y and shPR cells compared to control cells, confirming that the low PR levels of these cells affected the transcription of PR-target genes (Figure S2). In the same manner, the estradiol-mediated induction of *TFF1* gene transcription in control cells was strongly reduced in shPR and T47D-Y cells (Figure 1c), consistent with the reduced ERα level upon PR loss.

Finally, to test the role of PR in the maintenance of *ESR1* gene expression in different cellular backgrounds, we depleted the PR levels in two additional ER/PR positive cell lines, MCF7 and BT474, using the same short-hairpin RNA approach. Despite a moderate decrease of the PR levels in these cell lines, this slight decrease of PR was accompanied by a concomitant decrease of *ESR1* expression compared to control cells (Figure S1c,d), confirming the importance of PR levels in maintaining *ESR1* expression in different breast cancer cell lines.

### 2.2. PR Binds to the ESR1 Locus in Hormone-Free Breast Cancer Cells

To test whether PR directly regulates *ESR1* gene expression prior to hormone stimulation, we analyzed ChIP-seq data obtained with an antibody against PR in serum-starved T47D cells [14]. In the absence of hormones, PR appears to bind to two genomic regions within the *ESR1* locus, one located within the gene promoter (chromosome 6: 152,128,500-152,129,000 hg19) and another within the third intron (chromosome 6: 152,222,250-152,222,650 hg19) (Figure 2a). The specificity of these two hormone-free PR binding events was confirmed by ChIP-qPCR using T47D cells and PR-deficient cells (T47D-Y) or PR-depleted cells (shPR). PR bound to the promoter and the intronic regions in T47D cells, but not in T47D-Y cells (Figure 2b) or in shPR cells (Figure 2c). Strikingly, our analysis of previously published ChIP-seq experiments performed in the same conditions [2,15] revealed that the intronic sequence bound by PR in hormone-deprived T47D cells exhibited marks of active enhancers, including histone H3 mono-methylation on lysine 4 (H3K4me1) and DNase hypersensitivity (Figure 2a) [16].

**Figure 2.**
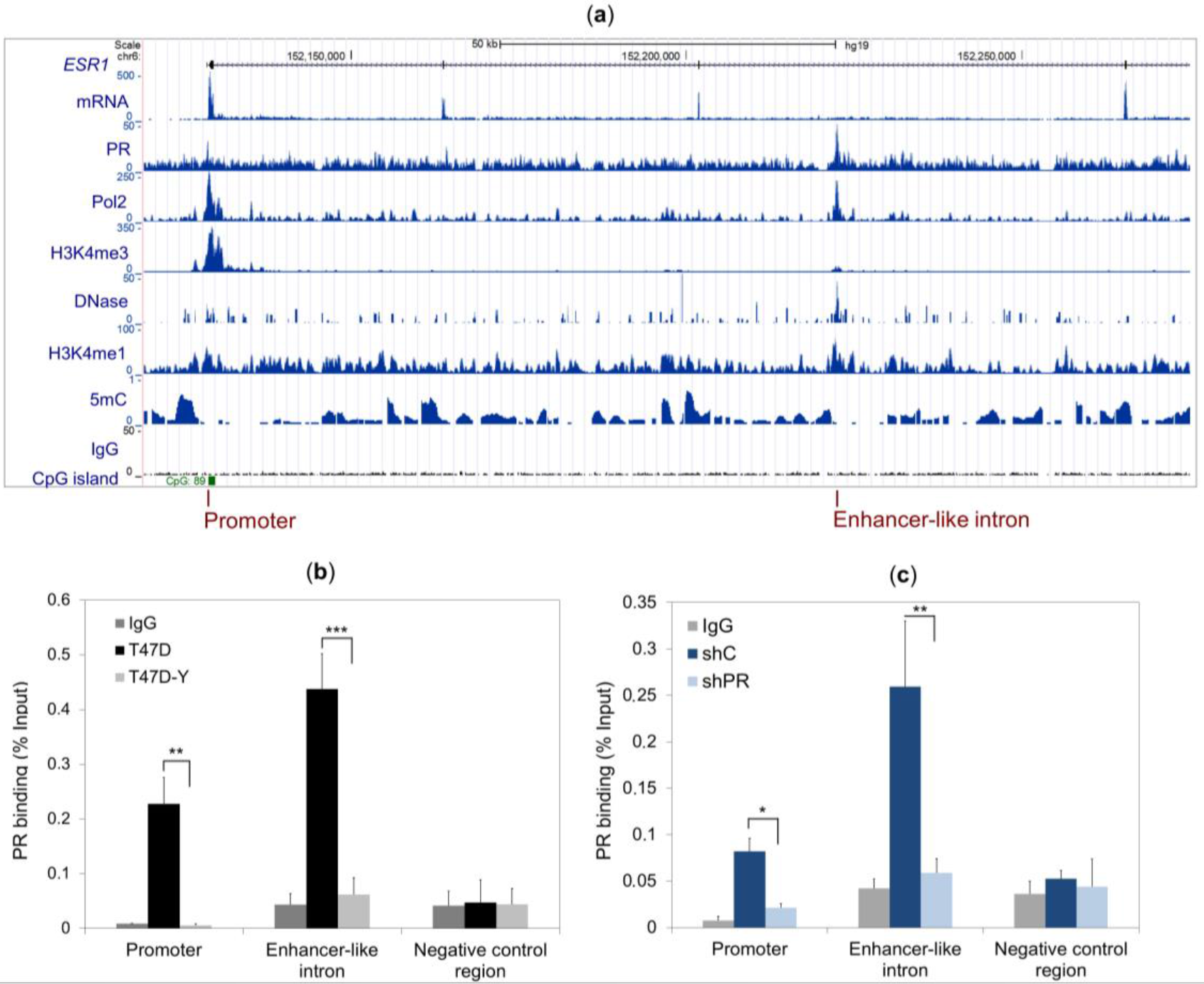
PR binds to the promoter and to an enhancer-like intron of the *ESR1* gene in hormone-deprived T47D breast cancer cells. (**a**) Screen shot from the UCSC genome browser showing the *ESR1* gene, the RNA reads and the ChIP-seq results from PR binding, with a peak at the gene promoter marked by polymerase 2 binding (Pol2), histone 3 trimethylated at lysine 4 (H3K4me3), low DNA methylation (5 mC) and a CpG island at the bottom. Another PR peak is found in an intronic region containing the classical enhancer epigenetic marks of the DNase hypersensitive site (DNase), histone 3 monomethylated at lysine 4 (H3K4me1) and low DNA methylation signal (5mC). The negative control immunoprecipitation is indicated by the IgG antibody. (**b**,**c**) The ChIP assay was performed with a specific antibody against PR or total rabbit IgG in T47D cells and PR-deficient cells (T47D-Y) (b) or PR-depleted cells (shPR, clone trcn0000010776) and control cells expressing a scrambled shRNA (shC) (c). Specific binding was assessed by quantitative PCR amplification of the *ESR1* gene promoter, an enhancer-like intronic sequence and a genomic region localized at the 3′-end of the enhancer-like intron (negative control region). Error bars represent the SD of three independent experiments. * *p* less than or equal to 0.05, ** *p* less than or equal to 0.01, *** *p* less than or equal to 0.005, unpaired twotailed Student’s *t*-test.

We also tested PR binding at the *ESR1* locus in MCF7 cells by ChIP-qPCR and found that in this case, the hormone-free PR also bound at the *ESR1* promoter, but not at the enhancer-like site in the third intron of the *ESR1* gene (Figure S3).

### 2.3. Rescue of PR Does Not Restore ESR1 Gene Expression in PR-Deficient Cells

To explore whether stable expression of PR restores *ESR1* gene expression, we stably-expressed PR in PR-deficient cells (T47D-Y +PR) and analyzed *ESR1* expression. Unexpectedly, *ESR1* expression remained significantly reduced at both transcript and protein levels after re-establishing PR levels (Figure 3a,b). Similarly, the estrogen-mediated induction of the *TFF1* gene remained reduced after PR rescue (Figure 3c). In contrast, the progestin-mediated gene transcription was significantly restored in PR-rescued cells (Figure S2), demonstrating the capability of re-expressed PR to mediate the PR-target gene transcription.

**Figure 3.**
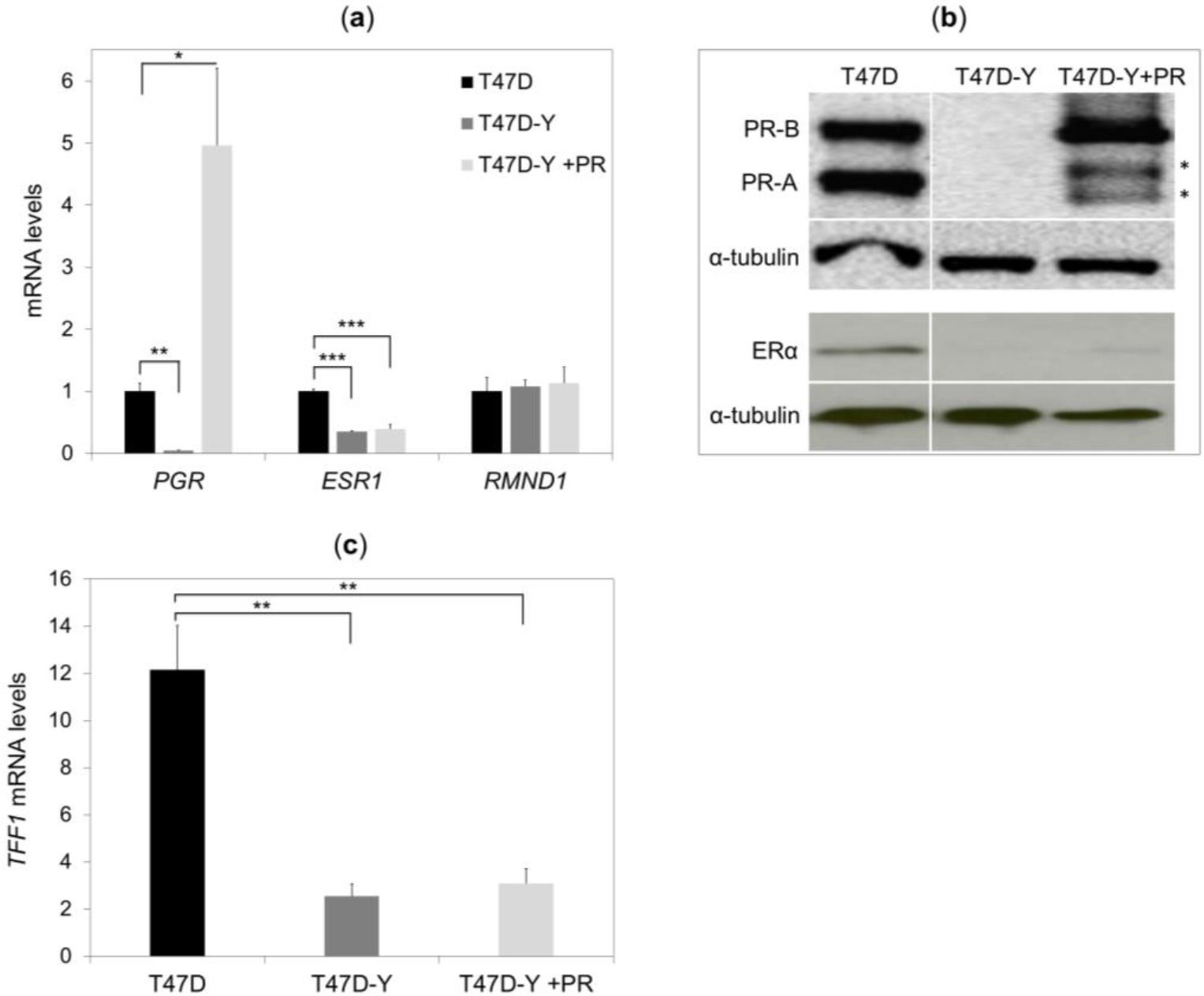
PR rescue of PR-deficient cells does not restore *ESR1* gene expression. (**a**) Gene-specific mRNA expression measured by quantitative RT-PCR in T47D control cells, PR-deficient cells (T47D-Y) and PR-rescue cells (T47D-Y + PR). The mRNA expression levels were normalized to *GAPDH* expression and are represented as values relative to the T47D cells. *PGR,* PR-encoding gene; *ESR1,* ER-encoding gene. Error bars represent the SD of three independent experiments. * *p* less than or equal to 0.05, ** *p* less than or equal to 0.01, *** *p* less than or equal to 0.005, unpaired two-tailed Student’s t-test. (**b**) Gene-specific protein levels measured by Western blotting in T47D control cells, T47D-Y and T47D-Y + PR cells. The lanes T47D and T47D-Y of this image are the same as in Figure 1b and are shown here for comparison with T47DY +PR. Blots are representative of three independent experiments. * indicates the degradation products of PR-B isoform. (**c**) PR rescue of PR-depleted cells does not restore the estrogen-mediated gene expression. T47D, PR-deficient cells (T47D-Y) and PR-rescue (T47D-Y + PR) cells were treated with estradiol (E2, 10 nM) or ethanol (vehicle) for 6 h, at which point, TFF1 mRNA expression levels were measured by quantitative RT-PCR. Gene-specific expression levels were normalized to *GAPDH* expression and are represented as values relative to the vehicle (E2/vehicle). Error bars represent the SD of three independent experiments. ** *p* less than or equal to 0.01, unpaired two-tailed Student’s *t*-test.

Together, these data demonstrate that PR re-expression alone is insufficient to restore *ESR1* gene expression to a level comparable to wild-type cells, suggesting that the *ESR1* gene is stably repressed through another mechanism once PR is absent in breast cancer cells.

### 2.4. Lack of PR Affects DNA Methylation at the ESR1 Promoter

DNA methylation at the *ESR1* promoter represents one of the main epigenetic mechanisms for stably repressing *ESR1* expression in breast cancers [17]. To explore whether PR loss affects the DNA methylation profile of the *ESR1* locus, we compared the DNA methylation pattern at the *ESR1* promoter and intronic PR-binding sites between T47D control cells, PR-deficient cells (T47D-Y) and PR-depleted cells (shPR) before and after PR rescue. Methylation of the *ESR1* promoter was strongly increased in PR-deficient cells and PR-depleted cells, and the higher DNA methylation levels of this genomic region persisted after PR rescue (T47D-Y +PR cells) (Figures 4 and S4). In contrast, the DNA methylation profile of the intronic-PR binding site was not increased, neither in PR-deficient, nor PR-depleted cells, and did not significantly change after PR rescue (Figure 4).

**Figure 4.**
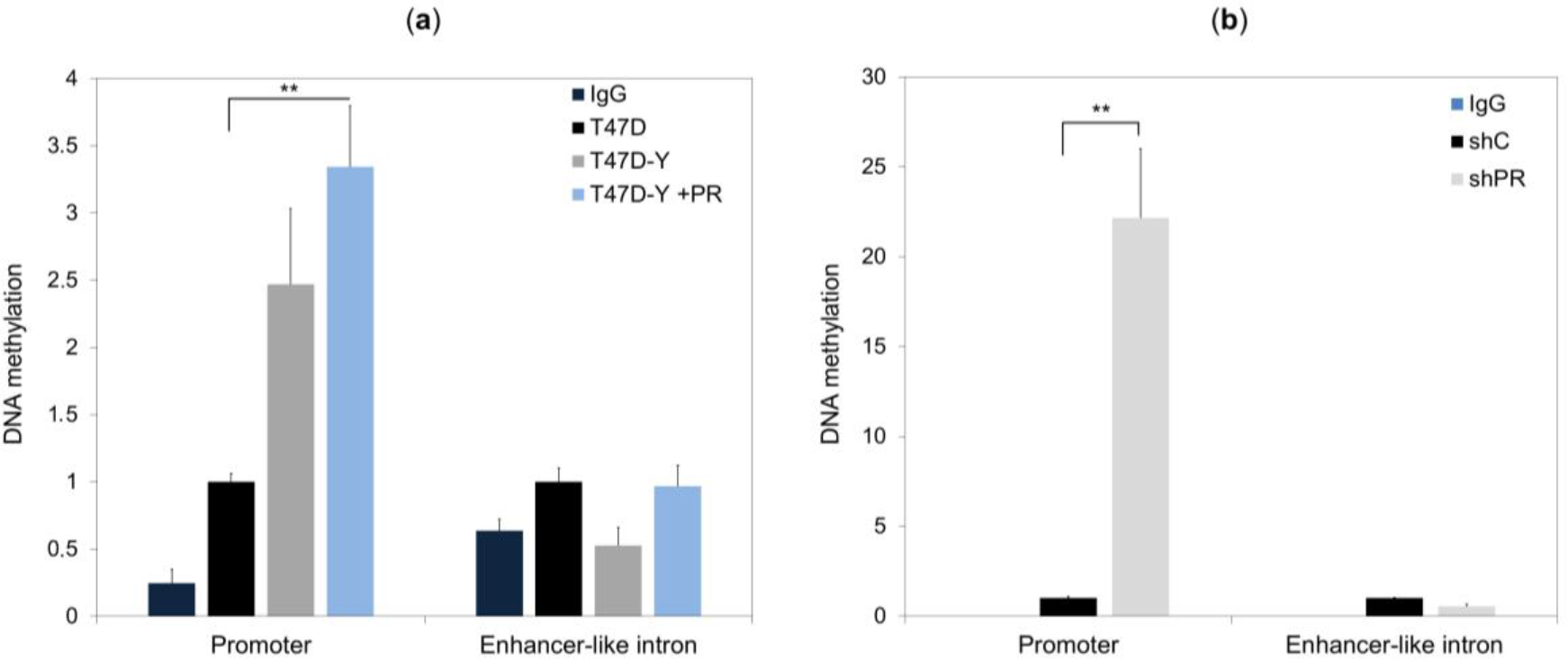
The loss of PR increases the DNA methylation level at the *ESR1* promoter. DNA methylation of the *ESR1* promoter and the enhancer-like intron was assessed by methylated DNA immunoprecipitation (MeDIP)-qPCR in T47D control cells, T47D-Y cells and T47D-Y cells with stable PR transfection (T47D-Y + PR) (**a**), or in T47D cells transduced with shRNA against *PR* (shPR; clone trcn0000010776) or scrambled shRNA (shC) (**b**). The results are represented as values relative to the control (T47D or shC). IgG, negative control for immunoprecipitation. Error bars represent the SD of three independent experiments. ** *p* less than or equal to 0.01, unpaired two-tailed Student’s *t*-test.

Consistent with these data, as previously demonstrated [18], the analysis of the TGCA breast invasive carcinoma (BRCA) dataset showed a clear difference in *ESR1* gene methylation levels when the patients were segregated based on the PR status. PR-negative breast carcinoma patients present higher methylation levels of the *ESR1* gene than PR-positive breast carcinomas. The gain of *ESR1* methylation in PR-negative breast carcinomas was stronger at the gene promoter rather than within the gene body (Figure S5).

### 2.5. DNA Methylation Impedes PR Binding to Hormone Responsive Elements

DNA methylation can directly affect the affinity of transcription factors towards their binding sites [19]. To check whether higher *ESR1* promoter methylation levels affect PR binding to this genomic region, we compared PR binding levels at the *ESR1* locus between T47D control cells, PR-deficient cells (T47D-Y) and PR-rescue cells (T47D-Y + PR) by the ChIP-qPCR assay. As described above, PR bound to the *ESR1* promoter and to an enhancer-like intronic sequence in control cells, whereas this binding was completely impaired in PR-deficient cells (Figure 2b). Rescue of PR in PR-deficient cells (T47D-Y +PR) completely restored PR binding at the low-methylated intronic sequence; however, it only partially restored PR binding at the highly-methylated promoter site, suggesting that hyper-methylation at the *ESR1* promoter impedes PR binding to this genomic region (Figure 5a). To test this hypothesis, we treated the PR-rescue cells (T47D-Y +PR) with the demethylating agent 5-azacytidine (5-azaC) or vehicle (control) and then compared the PR binding at the *ESR1* locus between control and 5-azaC-treated cells. The results showed that 5-azaC treatments significantly reduced DNA methylation levels at *ESR1* promoter (Figures 5b and S6) and had a tendency to increase PR binding levels at this genomic site (Figure 5c). In contrast, 5-azaC treatments did not affect the DNA methylation and the PR binding at the *ESR1* intronic region (Figure 5b,c).

**Figure 5.**
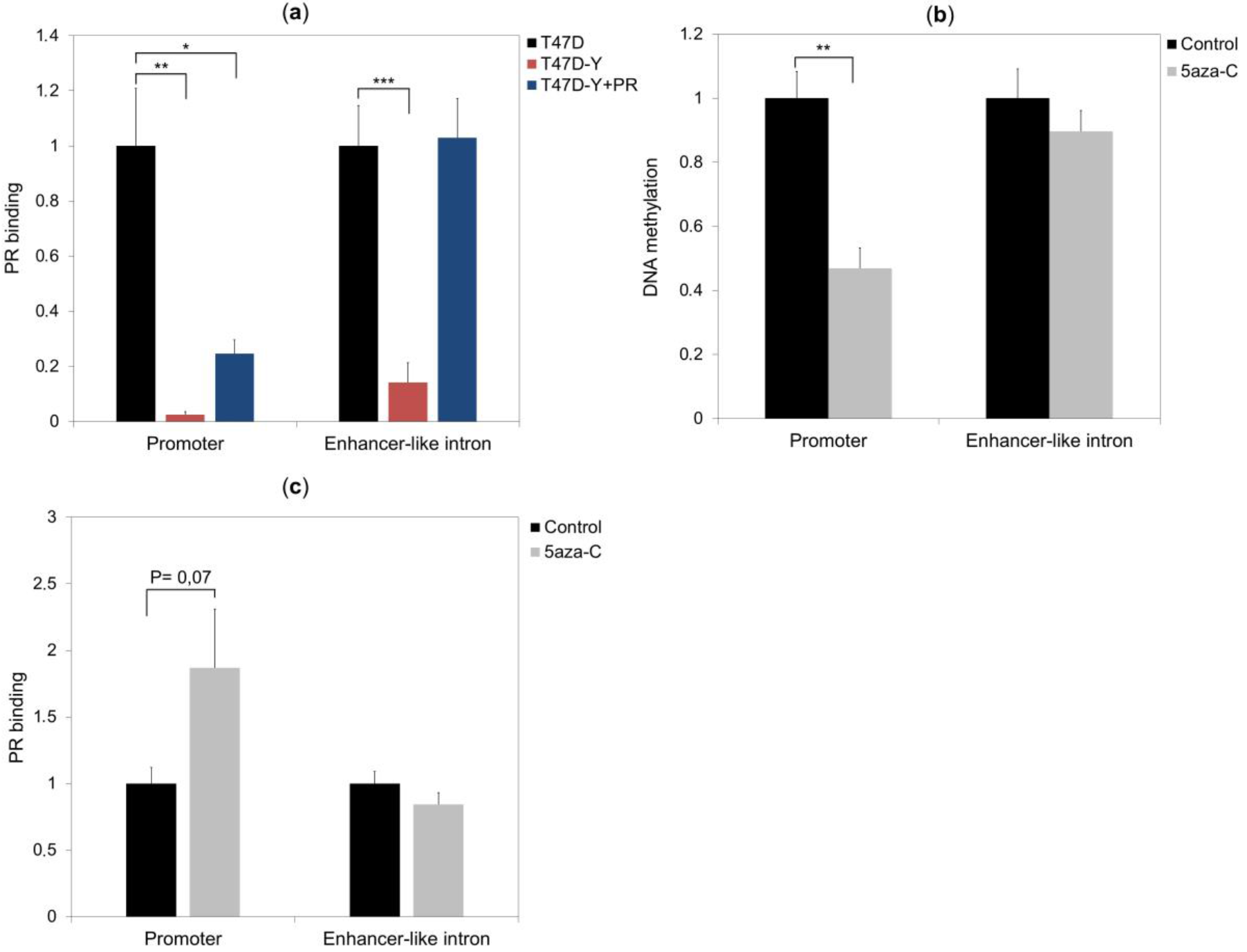
DNA methylation affects PR binding to the *ESR1* promoter. (**a**) The high-methylated *ESR1* promoter, in contrast to the low-methylated intronic sequence, was only partially bound by PR in PR-rescued cells (T47D-Y + PR). ChIP assays were performed with a specific antibody against PR. Specific binding was assessed by quantitative PCR amplification of the *ESR1* gene promoter and an enhancer-like intronic sequence in T47D control cells, PR-deficient cells (T47D-Y) and PR-rescued cells (T47D-Y + PR). Error bars represent the SD of three independent experiments. * *p* less than or equal to 0.05, ** *p* less than or equal to 0.01, *** *p* less than or equal to 0.005, unpaired two-tailed Student’s t-test. (**b**) The 5-azacytidine (5-azaC) demethylated *ESR1* promoter. DNA methylation analysis was performed by the MeDIP-qPCR assay using T47D-Y + PR cells treated with the demethylating agent 5-azaC (5 μM) or vehicle (control). The results are represented as fold change relative to the control. Error bars represent the SD of three independent experiments. ** *p* less than or equal to 0.01. (**c**) The demethylating agent 5-azaC increases PR binding at the *ESR1* promoter in PR-rescued cells (T47D-Y + PR). ChIP was performed as in (b) using T47D-Y + PR cells treated for 112 h with the demethylating agent 5-azaC (5 μM) or vehicle (control). Error bars represent the SD of three independent experiments. *p* = 0.07, unpaired two-tailed Student’s *t*-test.

The CpG island at the *ESR1* promoter contains a canonical progesterone-responsive elements (PRE) encompassing a CpG as well as six half palindromic PRE sites with one or two neighboring CpG (Figure 6a). To determine whether methylation of PRE affects the PR binding in vitro, we tested the PR binding to methylated or unmethylated forms of PRE oligonucleotides by the electrophoresis mobility shift assay (EMSA). We found that PR bound more efficiently to the unmethylated PRE oligonucleotides than to their methylated forms, especially when the PRE contained two CpG (Figure 6c), rather than one (Figure 6b). Further, in contrast to the methylated PRE, the unmethylated PRE oligonucleotide with two CpG was a high-affinity competitor in EMSA for an PRE oligonucleotide without CpG, which was previously shown to be a strong PR binding site (8) (Figure 6d). Interestingly, we observed that the CpG-containing PRE were bound by PR less efficiently than the PRE without CpGs (Figure 6b,c), suggesting that the presence of CpG, even if not methylated, reduces PR binding affinity. Finally, we analyzed the methylation of all genomic regions bound by hormone-free PR [14] and found that, despite having a higher CpG content than their flanking genomic regions, all the hormone-free PR binding sites had an overall lower level of methylation than their surrounding areas (Figure S7). These data suggest that high DNA methylation levels prevent PR binding, not only at the *ESR1* locus, but also at other PR binding sites.

**Figure 6.**
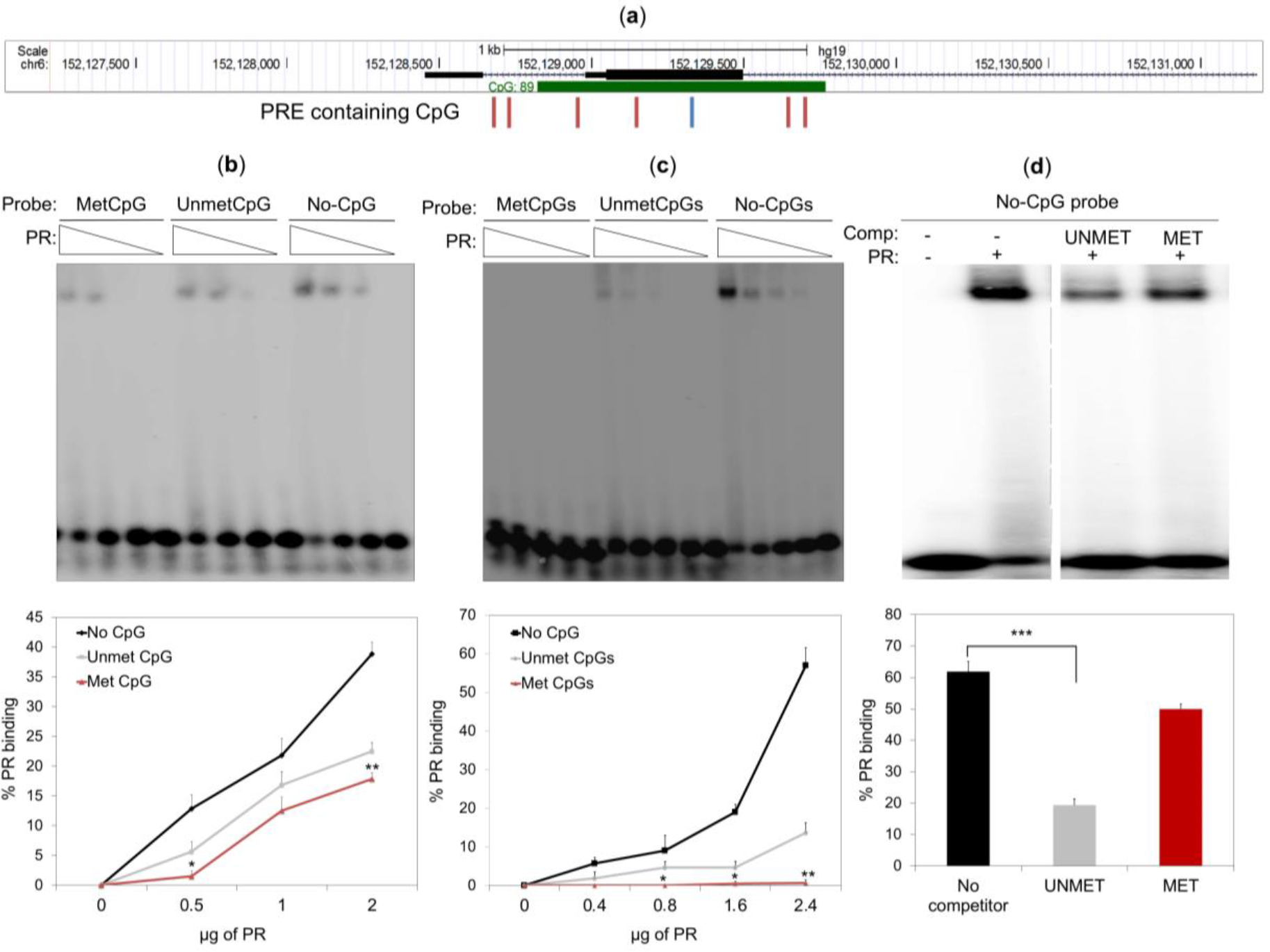
DNA methylation impedes PR binding to progesterone-responsive elements (PREs). (**a**) Screen shot from the UCSC genome browser showing the CpG island (CpG 89) at the *ESR1* promoter and the positions of the canonical PREs containing a CpG (blue line) and six half-palindromic PREs with one or two neighboring CpGs (red lines). (**b**,**c**) Electrophoretic-mobility shift assay using the indicated amount of purified human PR to capture the PRE with no CpG (ACAGTTTGT; no CpG), one methylated (MetCpG) or unmethylated CpG (UnmetCpG) (ACGGTTTGT) (**b**); two methylated (MetCpGs) or two unmethylated CpGs (UnmetCpGs) (ACGGTTCGT) (**c**). Quantification of the percentage of PR binding to different probes is shown in the lower part of the gel images. Error bars represent the SD of three independent experiments. * *p* less than or equal to 0.05, ** *p* less than or equal to 0.01, unpaired two-tailed Student’s *t*-test. (**d**) A double-stranded oligonucleotide probe with no CpG (ACAGTTTGT) was incubated with 2.4 μg of purified human PR and analyzed by PAGE either in the absence (−) or presence (+) of 100-fold excess of unlabeled oligonucleotides containing two unmethylated (UNMET) or two methylated (MET) CpGs (ACGGTTCGT). Error bars represent the SD of three independent experiments. *** *p* less than or equal to 0.005, unpaired two-tailed Student’s *t*-test. The dashed grey line indicates that a lane between the two samples was removed.

## 3. Discussion

The study of the role of PR in hormone-free breast cancer cells helps to clarify how these cells respond to external stimuli, including growth factors and ER modulators used for endocrine therapy. We show here that PR binds to the *ESR1* locus and is required to maintain the *ESR1* gene expression in hormone-free breast cancer cells. When the levels of PR are reduced, *ESR1* gene expression decreases in parallel with an increase of the DNA methylation level at the *ESR1* promoter, suggesting that unliganded PR maintains *ESR1* expression by preserving a low DNA methylation at the *ESR1* promoter. In line with these results, as previously demonstrated [18], our in silico analysis confirmed that PR-negative breast cancer patients present higher *ESR1* promoter methylation than PR-positive breast carcinomas. Consistently, demethylation of the *ESR1* promoter reactivates *ESR1* expression in ER-negative breast cancer cells [20].

The loss of PR did not affect DNA methylation of the *ESR1* intronic site, which in contrast to *ESR1* promoter lacks a CpG island, suggesting that PR selectively prevents methylation around CpG islands. However, whether the PR regulates *ESR1* promoter methylation in a direct or an indirect manner is not clear. Different molecular mechanisms activated by PR downregulation could be responsible for *ESR1* promoter hypermethylation in PR-depleted cells. In this context, it was suggested that DNA methylation can be a secondary event following gene silencing, rather than being a major event preceding it [21]. Thus, the reduced levels of *ESR1* transcription induced by PR depletion may drive to the gain of *ESR1* promoter methylation in PR-depleted cells.

In cells stably lacking PR expression, the stable re-expression of PR did not affect the hyper-methylation found at the *ESR1* promoter and was insufficient to reactivate *ESR1* gene expression. Moreover, rescue with PR completely restored the PR binding at the low-methylated intronic sequence, but only partially restored it at the highly methylated *ESR1* promoter site, suggesting that DNA methylation affects the PR binding to DNA. Consistently, the treatment of PR-rescued cells with the demethylating agent 5-azaC showed a tendency to increase PR binding at the *ESR1* promoter, but not at the low-methylated intronic site. Moreover, in vitro DNA binding experiments showed that PR preferentially bound unmethylated PRE oligonucleotides rather than their methylated counterpart, demonstrating that PR is a methylation-sensitive DNA binding protein. Taken together, these data demonstrate that the gain of DNA methylation at the *ESR1* promoter observed upon PR loss impedes PR binding to the *ESR1* promoter and maintains reduced *ESR1* transcription after PR rescue (Figure 7). Interestingly, there is a single human PR gene with two distinct promoter regions that encode two different isoforms, PR-A and PR-B. PR-B is the full-length protein that contains 933 amino acids, while PR-A lacks the first 164 amino acids. Although the two PR isoforms show high sequence similarity, they are functionally distinct transcriptional factors that can regulate the expression of a different subset of genes [22]. We showed that the depletion of both PR isoforms reduces the *ESR1* expression in T47D breast cancer cells; however, we cannot exclude that just one of the PR isoforms is required for the *ESR1* expression. Further analyses are required to clarify this possibility.

**Figure 7.**
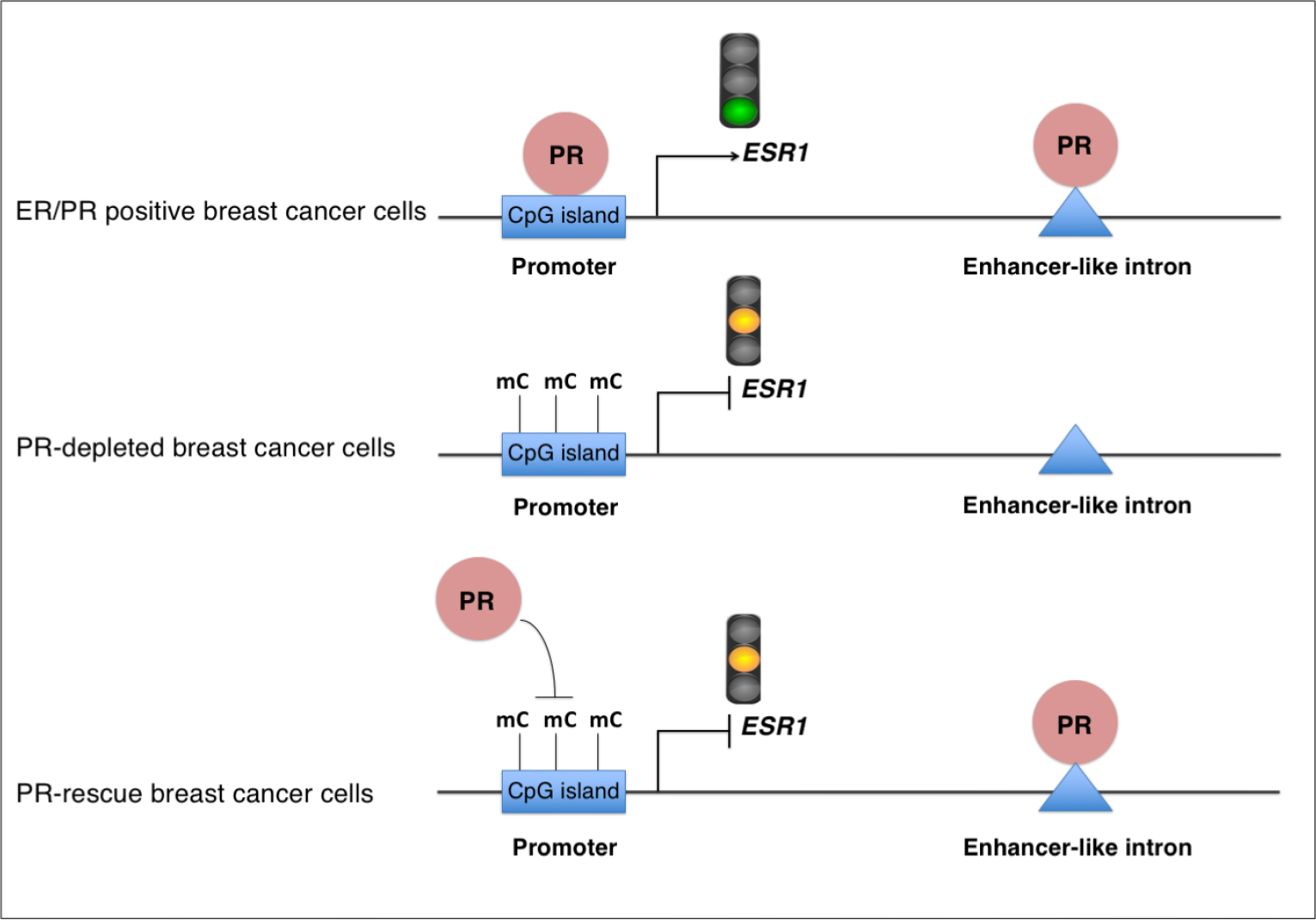
Model of the regulation of the *ESR1* gene expression and DNA methylation by PR in hormone-free breast cancer cells. In hormone-free ER+/PR+ breast cancer cells, PR binds to the low-methylated gene promoter, as well as to an enhancer-like intronic sequence of *ESR1.* PR binding at the gene promoter is required for maintaining *ESR1* transcription. In the absence of PR, DNA methylation (mC) increases at the ESR1 promoter, and ESR1 transcription is reduced. Re-expression of PR in PR-depleted cells leads to PR binding to the low-methylated enhancer-like intronic sequence, but the high level of DNA methylation (mC) at the ESR1 promoter impedes PR binding to this genomic region. Consequently, re-expression of PR in PR-depleted cells is insufficient to restore ESR1 expression.

In many cases, the transcriptional regulation of steroid target genes requires the action of regulatory sequences located far away from the promoters [23]. A significant fraction of these distal sequences engages in physical interactions with promoters, suggesting that they act as enhancers [22]. In this study, we showed that the PR binding site within the *ESR1* intronic sequence in T47D breast cancer cells exhibits the classical epigenetic marks found at active enhancer regions, including the monomethylation of lysine 4 of histone 3 (H3K4me1), a low DNA methylation and a DNase hypersensitivity [2,15]. This suggests that PR through its binding to the *ESR1* promoter and the enhancer-like intronic sequence could facilitate the interaction between these two genomic regions to enhance the *ESR1* transcription in T47D breast cancer cells.

The expression of ER and PR strongly affects breast cancer prognosis and response to endocrine therapy, with double-negative ER−/PR− breast cancers having worse clinical outcome than ER+/PR+ breast cancers [5]. Although ER is expressed in a large proportion of PR-positive breast cancers (ER+/PR+), a smaller percentage of patients express ER without PR (ER+/PR−) [4], demonstrating that a subgroup of ER+ breast cancers can still express the *ESR1* gene independently of the PR protein. However, the ER+/PR− breast cancers are less likely to respond to estrogen and selective ER modulator (SERM) therapy than breast cancers positive for both receptors (ER+/PR+) [5]. Interestingly, the different response to SERM therapy between these two groups of cancers was significant only in patients that were also negative for the epidermal growth factor 2 (HER2−), but not in HER2+ tumors, suggesting that the PR positively affects the response of breast cancers to endocrine therapy, especially when the HER2 signaling is inactivated [24]. Since the SERM treatments target ER, how the absence of PR affects this kind of therapy was unclear. Our finding that PR-deficient cells show lower ER levels compared to control cells suggests that the ER+/PR+ breast cancers are more likely to respond to SERM therapy because they could maintain the *ESR1* levels higher than breast cancers without PR protein. Future analyses are needed to clarify this hypothesis.

## 4. Materials and Methods

### 4.1. Cell Culture and Chemical Treatments

The T47D-MTVL (T47D) breast cancer cells used in this study have a stable integrated copy of the luciferase reporter gene driven by the MMTV promoter [14]. T47D, T47D-Y [11] and T47D-Y + PR [12] were routinely grown in RPMI 1640 medium and MCF7 and BT474 cells in DMEM medium. The T47D-Y + PR cell line was previously engineered to express the PR-B isoform [12]. The mediums were supplemented with 10% FBS, glutamine and standard antibiotics. For the experiments in the absence of hormones, cells were grown 48 h in medium without phenol red supplemented with 10% dextran-coated charcoal-treated FBS (DCC/FBS). In contrast to normal FBS, the DCC/FBS does not contain hormone-similar compounds that can activate PR activity. Moreover, since the cell cycle phase can influence PR activity [25], cells were then synchronized in G0/G1 by serum starvation for 16 h to avoid cell cycle phase variability. For the hormone treatments, cells were then induced with estradiol E2 (10 nM), progestin R5020 (10 nM) or vehicle (ethanol) for 6 h. For 5-azacytidine treatments, T47D-Y + PR cells were grown for 96 h (48 h using DMEM medium supplemented with 10% FBS and 48 h with DMEM without phenol red supplemented with 10% DCC/FBS) with 5 μM of 5-azacytidine (A3656, Sigma-Aldrich, St. Louis, MO, USA) or vehicle (1:1 acetic acid to water). Cells were finally synchronized in G0/G1 by 16 h of serum starvation before performing PR binding and DNA methylation analysis.

### 4.2. Lentivirus Preparation and Infection

HEK-293 cells were transfected with pVSV-G [26] and pCMVΔR8.91 [27], together with the pLKO.1-puro non-targeting vector (SHC001; Sigma-Aldrich, St. Louis, MO, USA) or pLKO.1-shRNA against PR using CaCl_2_ to permeabilize the cell membrane. Two different clones of PLKO.1-shRNA against PR have been used: clone trcn0000010776 and clone trcn0000003321 (SHCLND-NM_000926, Sigma-Aldrich, St. Louis, MO, USA). The viral particles containing the shRNA were collected 72 h after the transfection and used to infect breast cancer cells stably. Cell populations were finally selected with puromycin (1 μg/mL) and processed to quantify mRNA and protein expression.

### 4.3. Reverse Transcription and Quantitative PCR

Total RNA was isolated with the RNeasy extraction kit (Qiagen, Hilden, Germany). Complementary DNA (cDNA) was generated from 100 ng of total RNA with the First Strand cDNA Superscript II Synthesis kit (Invitrogen, Carlsbad, CA, USA; #11904018) and analyzed by quantitative PCR. Gene-specific expression was regularly normalized to *GAPDH* expression. Primers sequences are listed in Table S1.

### 4.4. Western Blotting

Cell lysates were resolved on SDS-polyacrylamide gels, and the proteins were transferred to Hybond-ECL nitrocellulose membranes (Amersham). Membranes were blocked with TBS-0.1% Tween 20 (TBS-T) with 5% of skimmed milk, incubated for 1 h at room temperature with a primary antibody (antibody against PR, sc-7208 from Santa Cruz Biotechnology; antibody against ERα, sc-543 from Santa Cruz Biotechnology; antibody against α-tubulin, T9026 from Sigma, St. Louis, MO, USA), diluted in TBS-T with 2.5% skimmed milk. After three washes with TBS-T, membranes were incubated for 45 min at room temperature with horseradish peroxidase-conjugated secondary antibodies (GE Healthcare, Chicago, IL, USA). Antibody binding was detected by chemiluminescence on an LAS-3000 image analyzer (Fuji PhotoFilm, Tokyo, Japan), and band intensity was quantified by the ImageJ tool.

### 4.5. Chromatin Immunoprecipitation

Chromatin immunoprecipitation (ChIP) assays were performed as described previously [28], with minor modifications. Cells were cross-linked in medium containing 1% formaldehyde for 10 min at 37 °C, and crosslinking was quenched with 125 mM glycine for 5 min at room temperature. After cells were lysed in hypotonic buffer, the nuclei were lysed with SDS-lysis buffer. Chromatin was sheared by sonication and incubated 16 h with 5 μg of antibody against progesterone receptor (PR, Santa Cruz Biotechnology, Dallas, TX, USA, sc-7208) or rabbit IgG (Cell Signaling, Danvers, MA, USA, #2729s). Immunocomplexes were recovered with protein A agarose bead slurry (Diagenode, Denville, NJ, USA, #C03020002) for 2 h with rotation and washed with ChIP buffer (Diagenode, Denville, NJ, USA, #K0249001) and Tris-EDTA buffer. For reversing the crosslinking, samples were incubated with proteinase K (10 mg/mL) at 65 °C for 16 h. DNA was purified and analyzed by quantitative PCR. Primer sequences are listed in Table S1.

For PR ChIP-seq analysis, the reads of the previously published PR ChIP-seq [14] were trimmed using Trimmomatic (Version 0.33) with the parameter values recommended by Bolger et al. [29]. The trimmed reads were aligned to the hg19 assembly version of the human genome using BWA (Version 0.7.12-r1039) [30]. The FASTA file containing the genome reference sequence of hg19 was downloaded from the UCSC Genome Browser discarding the random scaffolds and the alternative haplotypes from the reference sequence for the alignment [31]. The BWA-MEM algorithm and SAMtools (Version 1.2, using htslib 1.2.1) [32] were used to convert SAM files to BAM files and to sort them to retain only uniquely-aligned reads. The PR binding sites were identified with the MACS2 tool (Version 2.1.0.20150420) [33]. Peaks were additionally filtered until those remaining had a false discovery rate (FDR) *q*-value <10^−6^ and a 4-fold enrichment over the control sample (input), leaving 476 peaks for subsequent analyses.

### 4.6. DNA Methylation

The DNA methylation analyses were performed by the methylated DNA immunoprecipitation assay coupled with quantitative-PCR (MeDIP-qPCR) or high-throughput sequencing (MeDIP-seq). For MeDIP-qPCR, genomic DNA was randomly sheared by sonication to generate fragments between 300 and 700 bp. Sonicated DNA was denatured and then immunoprecipitated using an antibody against 5mC (Eurogentec; #BI-MECY-1000) or mouse IgG antibody, as previously described [34]. The immunocomplexes were recovered using 8 μL Dynabeads (M-280; Thermofisher, Waltham, Massachusetts), and the pull-down products were detected by quantitative-PCR. Primers sequences are listed in Table S1.

For MeDIP-seq, adaptors from the NEBNext Ultra DNA Library Prep Kit from Illumina were added to the fragmented DNA. Fragmented DNA was immunoprecipitated with antibody against 5mC as described above, and the amplified library was prepared using the NEBNext Ultra DNA Library Prep Kit for Illumina (E7370L) following the manufacturer’s instructions. Amplified libraries were sequenced, and reads were aligned with BowTie v1.1.2 using the reference human genome Version 19 (hg19), as previously described [35]. The mapped reads were filtered for duplicates and used to compute genome-wide reads-per-million (RPM) normalized signal tracks. The 5mC and CpG heat maps were generated using DeepTools (Version 2.2.0) [36] and BEDtools (Version v2.24.0) [37], and the matrix underlying the heatmaps was used to generate the 5mC and CpG average profiles. To test the significance of the overall reduction of 5mC methylation observed in the progesterone-receptor binding sites (PRBs), we calculated the average 5mC normalized read counts signal over each PRB and random regions resulting from shuffling the genomic coordinates of the PRBs, while keeping their sizes as in the true set of regions (this second step was repeated 1000 times to generate an empirical null distribution of 5mC methylation averaged values). The Mann-Whitney U-test was applied using the stats module of the Python’s SciPy library [38]. The DNA methylation data obtained by the MeDIP-seq method are available in the Gene Expression Omnibus (GEO) repository, Accession Number GSE107966.

For bisulfite-sequencing method, DNA was treated with sodium bisulfite using the EpiTect bisulfite kit (Qiagen, Hilden, Germany), and the *ESR1* promoter was amplified by two rounds of PCR using the oligonucleotide primers listed in Table S1. The PCR product was gel purified by the gel extraction kit (Qiagen, Hilden, Germany) and cloned in the pCR2.1 Topo TA cloning vector (Invitrogen, Carlsbad, CA, USA). Ten recombinant clones were then isolated using the GenElute plasmid miniprep kit (Sigma-Aldrich, St. Louis, MO, USA) and finally sequenced on an ABI DNA sequencer.

### 4.7. Electrophoresis Mobility-Shift Assay

Recombinant human PR (isoform B; PRB) was expressed in baculovirus and purified as previously described [39]. Radioactive double-stranded oligonucleotides containing the progesterone-responsive elements (PRE) were incubated with the indicated amounts of PR-B for 20 min at room temperature and analyzed in a 5% acrylamide-bisacrylamide electrophoresis gel. The radioactivity of the DNA-protein complex was then quantified by using the PhosphorImager and ImageQuant software (Molecular Dynamics). For the EMSA competition assay, a radioactive oligonucleotide without CpGs was first mixed with 100-fold of non-radioactive unmethylated or methylated probe containing two CpGs and then incubated with 2.4 μg of PR-B for 20 min at room temperature. DNA-protein complexes either in the absence or presence of unlabeled oligonucleotides were then analyzed as described above. Oligonucleotides sequences are listed in Table S1.

## 5. Conclusions

In this study, we demonstrate that PR binds to the low-methylated *ESR1* promoter and maintains both basal gene expression and the DNA methylation profile of the *ESR1* locus in hormone-free breast cancer cells. These data expand our understanding of the complex crosstalk between PR and ER and suggest that the analysis of DNA methylation of the *ESR1* promoter in breast cancer cells can help to design more appropriate targeted therapies for different types of breast cancer patients.

## Supplementary Materials

Figure S1: Depletion of PR reduces the *ESR1* expression in hormone-free breast cancer cells; Figure S2: PR depletion and PR rescue affect progestin-mediated gene transcription; Figure S3: PR binds at *ESR1* gene promoter in MCF7 breast cancer cells; Figure S4: The loss of PR increases DNA methylation at *ESR1* promoter; Figure S5: DNA methylation at the *ESR1* locus is higher in PR-negative than in PR-positive breast invasive cancers; Figure S6: The 5-azacytidine (5-azaC) demethylates the *ESR1* promoter; Figure S7: The PR binding sites are low methylated in hormone-deprived breast cancer cells. Table S1: Oligonucleotide sequences.

## Author Contributions

G.V., F.L.D., and M.B. designed the study and strategy of the project. G.V. and L.I.D.L. performed the experimental work. J.Q. provided the bioinformatics analysis of high throughput data. R.H.G.W. performed the analysis of the TGCA dataset. G.V. and F.L.D. discussed the results and wrote the paper. S.P. helped with the discussion of the results. All the authors contributed to editing the manuscript.

## Funding

We received funding from the Spanish Ministry of Economy and Competitiveness, Plan Nacional Project SAF 2013-42497-P; Centro de Excelencia Severo Ochoa 2013-2017; the Centre de Recerca de Catalunya (CERCA) Programme/Generalitat de Catalunya; G.V. has received funding from the Spanish Ministry of Economy and Competitiveness, “Juan de la Cierva Incorporation” fellowship (Ref. IJCI-2014-20723), the European Union Seventh Framework Programme (FP7/2007-2013) under Grant Agreement Number 299429 and the European Molecular Biology Organization (EMBO long-term fellowship ALTF 1106-2011, cofunded with the European Commission EMBOCOFUND2010, GA-2010-267146).

## Acknowledgments

We acknowledge support from Daniel Soronellas and Giancarlo Castellano for the DNA methylation mapping of MeDIP-sequencing analysis, Jofre Font for the PR purification, Enrique Vidal for revision of the statistical analysis and Silvina Nacht, Roberto Ferrari and Aldo Verde for their support.

## Conflicts of Interest

The authors declare no conflict of interest.

## References

1. Hilton, H.N.; Clarke, C.L.; Graham, J.D. Estrogen and progesterone signaling in the normal breast and its implications for cancer development. Mol. Cell. Endocrinol. 2018, 466, 2–14, pii:S0303-7207(17), 30433–1, doi:10.1016/j.mce.2017.08.011.

2. Ballaré, C.; Castellano, G.; Gaveglia, L.; Althammer, S.; González-Vallinas, J.; Eyras, E.; Le Dily, F.; Zaurin, R.; Soronellas, D.; Vicent, G.P.; et al. Nucleosome-Driven Transcription Factor Binding and Gene Regulation. Mol. Cell 2013, 49, 67–79, doi:10.1016/j.molcel.2012.10.019.

3. Wierer, M.; Verde, G.; Pisano, P.; Molina, H.; Font-Mateu, J.; Di Croce, L.; Beato, M. PLK1 signaling in breast cancer cells cooperates with estrogen receptor-dependent gene transcription. Cell Rep. 2013, 3, 2021–2032, doi:10.1016/j.celrep.2013.05.024.

4. Dunnwald, L.K.; Rossing, M.A.; Li, C.I. Hormone receptor status, tumor characteristics, and prognosis: A prospective cohort of breast cancer patients. Breast Cancer Res. 2007, 9, R6, doi:10.1186/bcr/bcr1639.

5. Cui, X.; Schiff, R.; Arpino, G.; Osborne, C.K.; Lee, A.V. Biology of progesterone receptor loss in breast cancer and its implications for endocrine therapy. J. Clin. Oncol. 2005, 23, 7721–7735, doi:10.1200/JC.O.2005.09.004.

6. Daniel, A.R.; Gaviglio, A.L.; Knutson, T.P.; Ostrander, J.H.; D’Assoro, A.B.; Ravindranathan, P.; Peng, Y.; Raj, G.V.; Yee, D.; Lange, C.A. Progesterone receptor-B enhances estrogen responsiveness of breast cancer cells via scaffolding PELP1- and estrogen receptor-containing transcription complexes. Oncogene 2015, 34, 506–515, doi:10.1038/onc.2013.579.

7. Encarnacion, C.A.; Ciocca, S.R.; McGuire, W.L.; Clark, G.M.; Fuqua, S.A.W.; Osborne, C.K. Measurement of steroid hormone receptors in breast cancer patients on tamoxifen. Breast Cancer Res. Treat. 1993, 26, 237–246.

8. Qiu, M.; Lange, C.A. MAP kinases couple multiple functions of human progesterone receptors: Degradation, transcriptional synergy, and nuclear association. J. Steroid Biochem. Mol. Biol. 2003, 85, 147–157, doi:10.1016/S0960-0760(03)00221-8.

9. Daniel, A.R.; Lange, C.A. Protein kinases mediate ligand-independent derepression of sumoylated progesterone receptors in breast cancer cells. Proc. Natl. Acad. Sci. USA 2009, 106, 14287–14292, doi:10.1073/pnas.0905118106.

10. Bendrat, K.; Fritz, P.; Müller, S.; Brockmöller, S.; Debus, A.; Friedrichs, K.; Lindner, C.; Brinkmann, F.; Heidemann, E.; Niendorf, A. Improved Risk Stratification for Breast Cancer Samples Based on the Expression Ratio of the Estrogen and Progesterone Receptor. Anticancer Res. 2016, 36, 3855–3863.

11. Sartorius, C.A.; Groshong, S.D.; Miller, L.A.; Powell, R.L.; Tung, L.; Takimoto, G.S.; Horwitz, K.B. New T47D breast cancer cell lines for the independent study of progesterone B- and A-receptors: Only antiprogestin-occupied B-receptors are switched to transcriptional agonists by cAMP. Cancer Res. 1994, 54, 3868–3877.

12. Quiles, I.; Millán-Ariño, L.; Subtil-Rodríguez, A.; Miñana, B.; Spinedi, N.; Ballaré, C.; Beato, M.; Jordan, A. Mutational analysis of progesterone receptor functional domains in stable cell lines delineates sets of genes regulated by different mechanisms. Mol. Endocrinol. 2009, 23, 809–826, doi:10.1210/me.2008-0454.

13. Alexander, I.E.; Shine, J.; Sutherland, R.L. Progestin regulation of estrogen receptor messenger RNA in human breast cancer cells. Mol. Endocrinol. 1990, 6, 821–828, doi:10.1210/mend-4-6-821.

14. Vicent, G.P.; Nacht, A.S.; Zaurin, R.; Font-Mateu, J.; Soronellas, D.; Le Dily F.; Reyes, D.; Beato, M. Unliganded progesterone receptor mediated targeting of an RNA-containing repressive complex silences a subset of hormone-inducible genes. Genes Dev. 2013, 27, 1179–1197, doi:10.1101/gad.215293.113.

15. Le Dily, F.; Baù, D.; Pohl, A.; Vicent, G.P.; Serra, F.; Soronellas, D.; Castellano, G.; Wright, R.H.; Ballaré, C.; Filion, G.; et al. Distinct structural transitions of chromatin topological domains correlate with coordinated hormone-induced gene regulation. Genes Dev. 2014, 28, 2151–2162, doi:10.1101/gad.241422.114.

16. Calo, E.; Wysocka, J. Modification of enhancer chromatin: What, How, and Why? Mol. Cell 2013, 49, 825–837, doi:10.1016/j.molcel.2013.01.038.

17. Martínez-Galán, J.; Torres-Torres, B.; Núñez, M.I.; López-Peñalver, J.; Del Moral, R.; Ruiz De Almodóvar, J.M.; Menjón, S.; Concha, A.; Chamorro, C.; Ríos, S.; et al. ESR1 gene promoter region methylation in free circulating DNA and its correlation with estrogen receptor protein expression in tumor tissue in breast cancer patients. BMC Cancer 2014, 14, 59, doi:10.1186/1471-2407-14-59.

18. Widschwendter, M.; Siegmund, K.D.; Müller, H.M.; Fiegl, H.; Marth, C.; Müller-Holzner, E.; Jones, P.A.; Laird, P.W. Association of breast cancer DNA methylation profiles with hormone receptor status and response to tamoxifen. Cancer Res. 2004, 64, 3807–3813, doi:10.1158/0008-5472.CAN-03-3852.

19. Yin, Y.; Morgunova, E.; Jolma, A.; Kaasinen, E.; Sahu, B.; Khund-Sayeed, S.; Das, P.K.; Kivioja, T.; Dave, K.; Zhong, F.; et al. Impact of cytosine methylation on DNA binding specificities of human transcription factors. Science 2017, 356, pii:eaaj2239, doi:10.1126/science.aaj2239.

20. Ferguson, A.T.; Lapidus, R.G.; Baylin, S.B.; Davidson, N.E. Demethylation of the estrogen receptor gene in estrogen receptor-negative breast cancer cells can reactivate estrogen receptor gene expression. Cancer Res. 1995, 55, 2279–2283.

21. Jones, P.A. Functions of DNA methylation: Islands, start sites, gene bodies and beyond. Nat. Rev. Genet. 2012, 13, 484–492, doi:10.1038/nrg3230.

22. Richer, J.K.; Jacobsen, B.M.; Manning, N.G.; Abel, M.G.; Wolf, D.M.; Horwitz, K.B. Differential gene regulation by the two progesterone receptor isoforms in human breast cancer cells. J. Biol. Chem. 2002, 277, 5209–5218, doi:10.1074/jbc.M110090200.

23. Carroll, J.S.; Liu, X.S.; Brodsky, A.S.; Li, W.; Meyer, C.A.; Szary, A.J.; Eeckhoute, J.; Shao, W.; Hestermann, E.V.; Geistlinger, T.R.; et al. Chromosome-wide mapping of estrogen receptor binding reveals long-range regulation requiring the forkhead protein FoxA1. Cell 2005, 122, 33–43, doi:10.1016/j.cell.2005.05.008.

24. Bae, S.Y.; Kim, S.; Lee, J.H.; Lee, H.C.; Lee, S.K.; Kil, W.H.; Kim, S.W.; Lee, J.E.; Nam, S.J. Poor prognosis of single hormone receptor-positive breast cancer: similar outcome as triple-negative breast cancer. BMC Cancer 2015, 15, 138, doi:10.1186/s12885-015-1121-4.

25. Dressing, G.E.; Knutson, T.P.; Schiewer, M.J.; Daniel, A.R.; Hagan, C.R.; Diep, C.H.; Knudsen, K.E.; Lange, C.A. Progesterone receptor-cyclin D1 complexes induce cell cycle-dependent transcriptional programs in breast cancer cells. Mol. Endocrinol. 2014, 28, 442–457, doi:10.1210/me.2013-1196.

26. Stewart, S.A.; Dykxhoorn, D.M.; Palliser, D.; Mizuno, H.; Yu, E.Y.; An, D.S.; Sabatini, D.M.; Chen, I.S.; Hahn, W.C.; Sharp, P.A.; et al. Lentivirus-delivered stable gene silencing by RNAi in primary cells. RNA 2003, 9, 493–501.

27. Zufferey, R.; Nagy, D.; Mandel, R.J.; Naldini, L.; Trono, D. Multiply attenuated lentiviral vector achieves efficient gene delivery in vivo. Nat. Biotechnol. 1997, 15, 871–875, doi:10.1038/nbt0997-871.

28. Strutt, H.; Paro, R. Mapping DNA target sites of chromatin proteins in vivo by formaldehyde crosslinking. Methods Mol. Biol. 1999, 119, 455–467, doi:10.1385/1-59259-681-9:455.

29. Bolger, A.M.; Lohse, M.; Usadel, B. Trimmomatic: A flexible trimmer for Illumina sequence data. Bioinformatics 2014, 30, 2114–2120, doi:10.1093/bioinformatics/btu1701.

30. Li, H.; Durbin, R. Fast and accurate short read alignment with Burrows-Wheeler transform. Bioinformatics 2009, 25, 1754–1760, doi:10.1093/bioinformatics/btp324.

31. Karolchik, D.; Barber, G.P.; Casper, J.; Clawson, H.; Cline, M.S.; Diekhans, M.; Dreszer, T.R.; Fujita, P.A.; Guruvadoo, L.; Haeussler, M.; et al. The UCSC Genome Browser database: 2014 update. Nucleic Acids Res. 2014, 42, D764–D770, doi:10.1093/nar/gkt1168.

32. Li, H.; Handsaker, B.; Wysoker, A.; Fennell, T.; Ruan, J.; Homer, N.; Marth, G.; Abecasis, G.; Durbin, R. 1000 Genome Project Data Processing Subgroup. The sequence Alignment/Map format and SAMtools. Bioinformatics 2009, 25, 2078–2079, doi:10.1093/bioinformatics/btp352.

33. Zhang, Y.; Liu, T.; Meyer, C.A.; Eeckhoute, J.; Johnson, D.S.; Bernstein, B.E.; Nusbaum, C.; Myers, R.M.; Brown, M.; Li, W.; et al. Model-based analysis of ChIP-Seq (MACS). Genome Biol. 2008, 9, R137, doi:10.1186/gb-2008-9-9-r137.

34. Mohn, F.; Weber, M.; Schübeler, D.; Roloff, T.C. Methylated DNA immunoprecipitation (MeDIP). Methods Mol. Biol. 2009, 507, 55–64, doi:10.1007/978-1-59745-522-0_5.

35. Langmead, B.; Trapnell, C.; Pop, M.; Salzberg, S.L. Ultrafast and memory-efficient alignment of short DNA sequences to the human genome. Genome Biol. 2009, 10, R25, doi:10.1186/gb-2009-10-3-r25.

36. Ramírez, F.; Ryan, D.P.; Grüning, B.; Bhardwaj, V.; Kilpert, F.; Richter, A.S.; Heyne, S.; Dündar, F.; Manke, T. deepTools2: A next generation web server for deep-sequencing data analysis. Nucleic Acids Res. 2016, 44, 160–165, doi:10.1093/nar/gkw257.

37. Quinlan, A.R. BEDTools: The swiss-army tool for genome feature analysis. Curr. Protoc. Bioinform. 2014, 47, doi:10.1002/0471250953.bi1112s47.

38. Oliphant, T.E. Python for scientific computing. Comput. Sci. Eng. 2007, 9, 10–20, doi:10.1109/MCS.E.2007.58.

39. Di Croce, L.; Koop, R.; Venditti, P.; Westphal, H.M.; Nightingale, K.P.; Corona, D.F.; Becker, P.B.; Beato, M. Two-step synergism between the progesterone receptor and the DNA-binding domain of nuclear factor 1 on MMTV minichromosomes. Mol. Cell 1999, 4, 45–54, doi:10.1016/S1097-2765(00)80186-0.

